# Loss of critical developmental and human disease-causing genes in 58 mammals

**DOI:** 10.1101/819169

**Authors:** Yatish Turakhia, Heidi I. Chen, Amir Marcovitz, Gill Bejerano

## Abstract

Gene losses provide an insightful route for studying the morphological and physiological adaptations of species, but their discovery is challenging. Existing genome annotation tools and protein databases focus on annotating intact genes and do not attempt to distinguish nonfunctional genes from genes missing annotation due to sequencing and assembly artifacts. Previous attempts to annotate gene losses have required significant manual curation, which hampers their scalability for the ever-increasing deluge of newly sequenced genomes. Using extreme sequence erosion (deletion and non-synonymous substitution) as an unambiguous signature of loss, we developed an automated approach for detecting high-confidence protein-coding gene loss events across a species tree. Our approach relies solely on gene annotation in a single reference genome, raw assemblies for the remaining species to analyze, and the associated phylogenetic tree for all organisms involved. Using the hg38 human assembly as a reference, we discovered over 500 unique human genes affected by such high-confidence erosion events in different clades across 58 mammals. While most of these events likely have benign consequences, we also found dozens of clade-specific gene losses that result in early lethality in outgroup mammals or are associated with severe congenital diseases in humans. Our discoveries yield intriguing potential for translational medical genetics and for evolutionary biology, and our approach is readily applicable to large-scale genome sequencing efforts across the tree of life.

## Introduction

The placental mammal radiation exhibits tremendous phenotypic diversity [1]. These myriad forms and functions arose from modification of inherited traits from one generation to the next. An extreme case of this “descent with modification” involves disappearance (either sudden or gradual) of ancestral phenotypes, caused by one or more function-inactivating mutations. If it fixes in the species, this functional inactivation will be accompanied by relaxation of purifying selection on all genomic regions encoding the trait, allowing erosion of the sequence. When a gene loses its functional role this way, the result can be a so-called “gene loss” – a nonfunctional, dead gene [2].

Previous studies have identified a few intriguing examples of gene losses that are associated with phenotypic traits. For example, the enzyme gene *GULO* is inactivated in independent mammalian lineages that consequently lost the ability to synthesize Vitamin C, and must instead have a constant supply of it in their diet [3, 4]; multiple visual system genes are no longer functional in subterranean mammals which live primarily in darkness [5, 6]; taste receptor and other genes are lost in fully aquatic cetaceans [7–9]; renal transporter genes *URAT1*, *GLUT9* and *OAT1* are dead in fruit-eating bats, which could have facilitated their frugivorous diet [10]; the immune genes *MX1* and *MX2* are eroded in toothed whales, possibly making them more susceptible to certain viral pathogens [11]; and the loss of *PON1* in several marine mammals may confer increased vulnerability to agricultural pesticide pollution [12]. Systematic annotation of gene losses across species may therefore not only reveal fascinating evolutionary events and genotype-phenotype relationships, but could also point us to “natural knockout” models for human pathologies that reveal compensating molecular pathways in species missing otherwise-indispensable genes [2, 3, 13].

Manual annotation of pseudogenized genes, such as by ENCODE-HAVANA [14], while accurate, has been applied to only a handful of species (human, mouse, rat and zebrafish) and is not scalable for hundreds to thousands of newly sequenced species [15]. Automating the discovery of gene losses is scalable but challenging. Recently, Sharma et al. [10] developed an automated method using whole-genome multiple-sequence alignment to identify genes with inactivating mutations. However, their overly inclusive approach predicted hundreds of gene losses per species across 62 mammals, of which only 21 total were reported, presumably because manual inspection found many loss predictions to be spurious. Similar technique was used to further semi-manually annotate 85 gene losses in cetaceans [16]. Other computational approaches for gene annotation, like the Ensembl pipeline [17], combine *ab initio* predictions with protein sequence evidence [18] and with gene models inferred from RNA-seq data. However, they focus only on annotating *intact* genes and do not attempt to distinguish decaying genes from those that only confoundingly appear so, for instance, as a result of sequencing gaps and alignment errors. By design, these tools are thus limited in their ability to identify gene loss events.

We developed an improved, conservative approach for identifying gene loss events and applied it to 58 mammalian genomes, using human assembly hg38 as the reference. We hypothesized that the most accurate strategy for declaring an ancestral gene function to be lost is to demonstrate that it is highly sequence-eroded compared to other genes in the same genome. While it is impossible to guarantee that no gene transcript is expressed from the orthologous locus (especially if its promoter remains intact), the erosion of gene sequence implies that the gene does not maintain its original function. To that end, we used pairwise whole-genome alignments to locate the orthologs of human protein-coding genes in each of the 58 query species by applying a novel synteny-aware mapping procedure. We employed a Mahalanobis distance-based model to identify the most highly mutated orthologs in each genome in a way that normalizes for the baseline extent of sequence divergence between this species and the reference (human). We considered these per-genome predictions to be *likely eroded* and functionally dead orthologs in the species represented by the assembly. Lastly, we applied a stringent phylogenetic filter, requiring that all final candidate erosions were supported by observations from at least two closely related species and affected ancestral genes – in order to minimize confounders due to assembly artefacts, private (individual-specific) mutations, or reference genome biases. Though very conservative, our approach uncovered hundreds of *<u>hi</u>gh-<u>conf</u>idence ortholog <u>erosion</u>* (*hiconfErosion*) events, affecting over 50 mammals, including some very surprising gene losses with intriguing evolutionary and biomedical implications. Importantly, our method was designed to allow easy addition of genomes to the analysis, such as the those from countless newly sequenced mammals, birds [19], vertebrates [20], insects [21] and more.

## Results

### Pre-existing gene annotation methods fail to identify and determine the functional state for thousands of orthologs

Of the 59 mammalian genomes (including the human reference) that we used in our study, we found Ensembl gene predictions (release 86) for only 34 genomes. The number of protein-coding gene annotations varied widely (Figure 1A), from 11,765 annotations in alpaca to 22,285 annotations in human. When plotting the number of Ensembl predicted genes against the assembly N50, it is clear that the annotation completeness is artifactually dependent on assembly contiguity or lack thereof (Figure 1A). For instance, absence of the critical chordate development gene *PAX6* in the dog annotation set is likely due to a sequencing gap in the genome assembly (Figure 1B). Similarly, while *TP53* and multiple adjacent genes are absent from the alpaca annotation set, its genome includes a region with high sequence similarity to the human *TP53* ortholog (Figure 1C), implying a potentially missed prediction. We have observed that pseudogene annotation pipelines are also incomplete – for example, manual curation of mouse pseudogenes by ENCODE-HAVANA has been on-going for several years, but the list is still not available online (https://www.gencodegenes.org/). Based on these observations, we devised an improved approach for identifying a high-confidence, conservative subset of gene loss events across the 58 placental mammals, using human as the reference.

**Figure 1.**
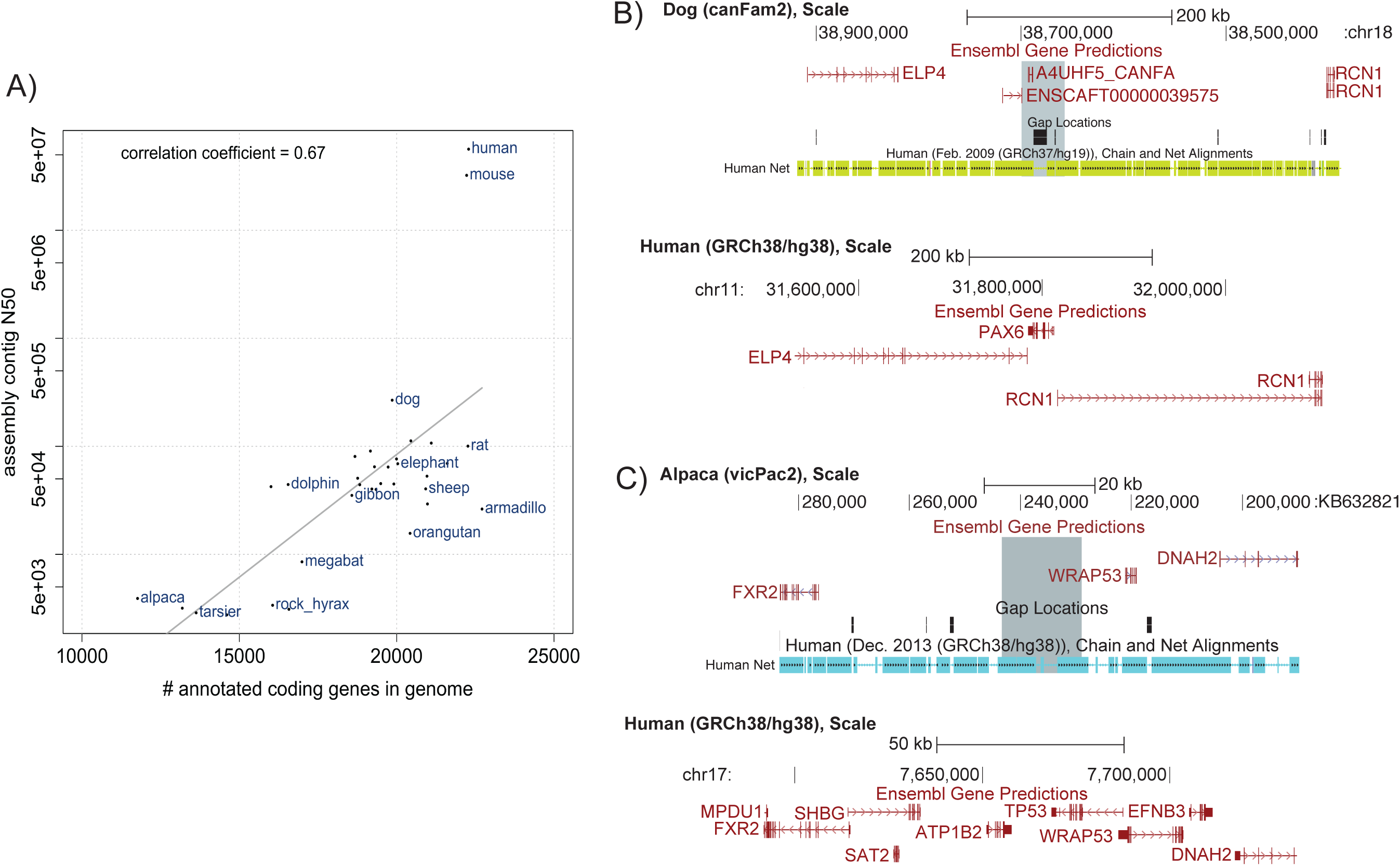
Incomplete coding gene annotations in genomic databases. A) Of the 59 mammalian genomes used in this study, 34 have Ensembl (release 86) gene annotations. The number of annotations per genome appears to artifactually increase with assembly quality (as measured by N50 contig length), with a correlation coefficient of 0.67, saturating at some point beyond N50≈5e+5 for highly contiguous assemblies (such as human and mouse). B) Top: *PAX6* Ensembl gene model absence (gray highlight) in the dog genome assembly due to a sequencing gap. The genome browser view shows the minus strand. Bottom: The orthologous position of *PAX6* (and flanking genes) in the human genome. C) Top: *TP53* Ensembl gene model absence (gray highlight) in the alpaca genome within a conserved region. The genome browser view shows the minus strand. Bottom: The orthologous position of *TP53* (and flanking genes) in the human genome.

### Mapping gene orthologs across mammalian genomes

To identify *ab initio* orthologs of human genes across the 58 query species, we used whole-genome pairwise alignments (i.e. Jim Kent’s BLASTZ-based *chains* [22]) to map genes from human coordinate space to each of the other genomes. Chains identify conserved sequences not only in coding regions (covering 2% of the human genome) but also in non-coding, gene-regulatory regions (covering 5-10% of the human genome) in which the genes are interspersed. As such, chains facilitate accurately determining genomic orthology between species. Using these 58 pairwise alignment chains anchored to the human reference, we determined the single locus in the aligning (query) genome that exhibited the greatest sequence similarity and surrounding synteny with each human protein-coding gene. Specifically, each successfully mapped gene was (i) positioned within a chain that had alignment and gene-in-synteny scores sufficiently higher than the next-highest-scoring (likely paralogous) chain and (ii) did not overlap in genomic space with any other gene mappings proposed in the previous step (see Methods, Figure 2A,B). In total, our mapping procedure identified 11,155 – 17,690 orthologs per species (median: 16,782; Figure 3A) across the 58 non-human mammals considered.

**Figure 2.**
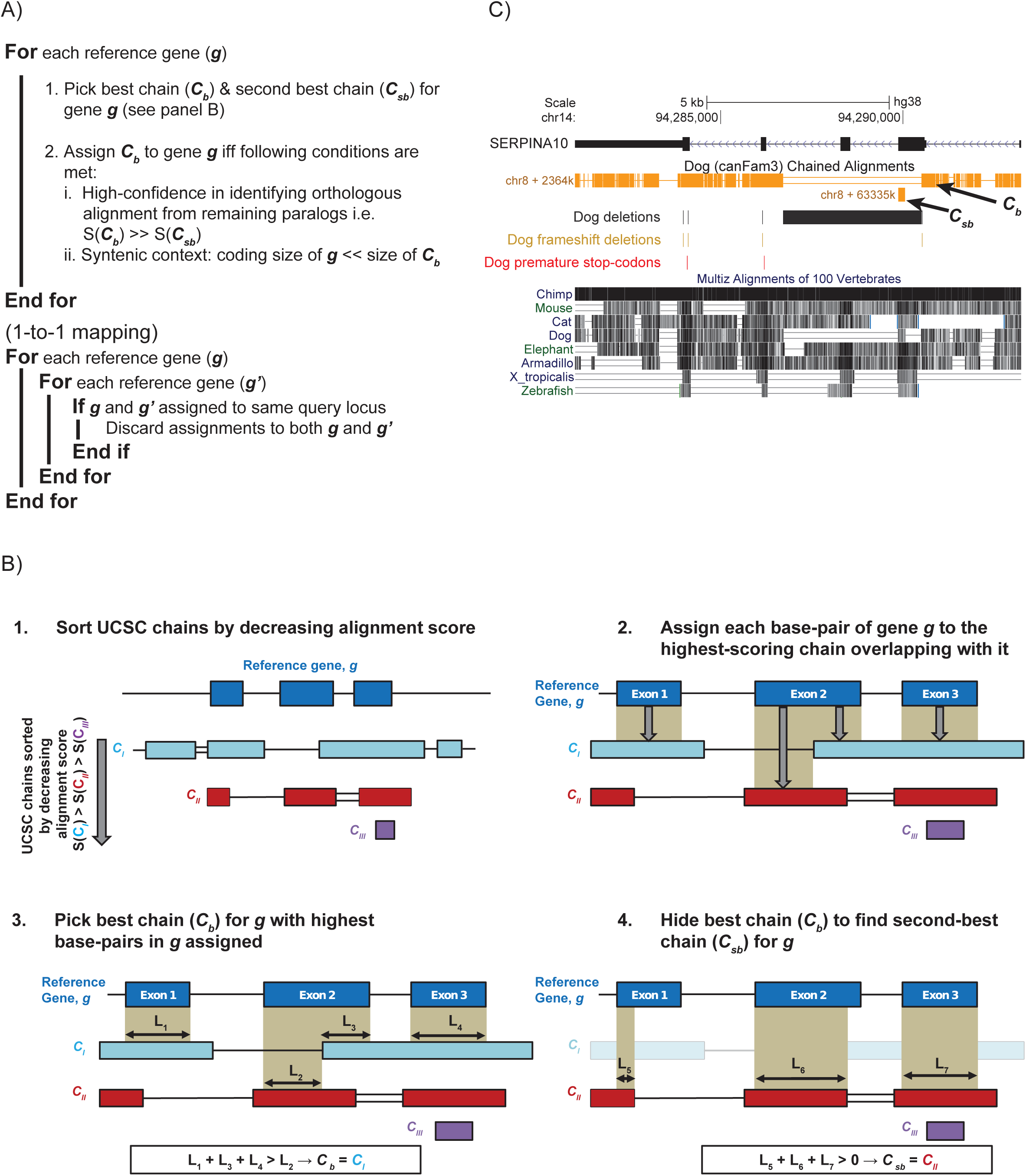
An overview of the chain-picking method to map genes from a reference genome to a locus in query species genome. A) We designed a conservative gene-by-gene scanning approach that maps genes from a reference genome (human, GRCh38) to their orthologous locations in target mammalian genomes based on pairwise genome chain alignments. For each reference gene, our approach first picks two chains: the *best chain* (*C_b_*) and the *second-best chain*, *C_sb_* (panel B). The chain *C_b_* is then assigned to a gene *g* if three additional conditions (as shown, also see Methods) are satisfied. B) Detailed illustration of the procedure to pick the *best-chain* (*C_b_*) and the *second-best chain* (*C_sb_*) for gene *g*. In the case of multiple chains at a locus (gene, *g*), our procedure first assigns each base of *g* to the highest-scoring chain overlapping with the base-pair, and then identifies the chain (*C_b_*) to which the most bases of *g* were assigned. The procedure is repeated to identify another chain (*C_sb_*) after *C_b_* is hidden. C) *SERPINA10* is a known loss of function gene in dogs [23], and hence lacks any annotations or orthology mapping from human to dog. This pairwise, chain-based approach enables precise mapping of an intact gene from a reference assembly to even sequence-eroded loci in another species.

**Figure 3.**
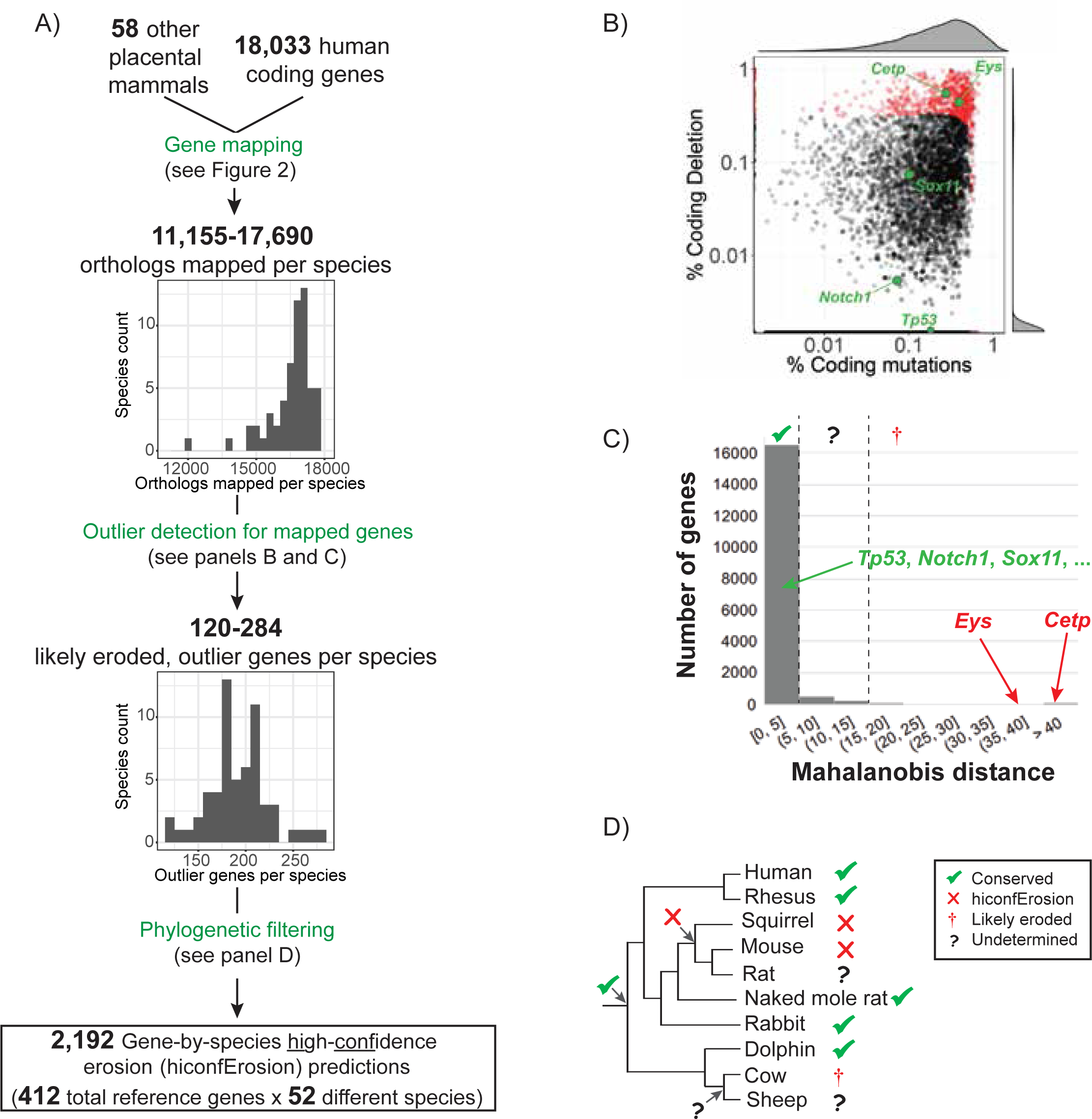
A pipeline to call hiconfErosions in genes across mammalian lineages. A) Using pairwise chained alignments, we determined an orthologous location for human coding genes in each genome. Then we computed the amount of coding deletion and amino acid substitution to find, for each each genome, the outlier genes among all the genes that were mapped (i.e., likely eroding genes). B) Scatter plot showing the distribution of the mouse (mm10) genes along the two features (% coding deletion and % coding substitution) for calculating Mahalanobis Distance (*MD*). The histogram along each axis shows the distribution of points when projected on the respective axis. (Per-species) likely eroded genes (in red, such as known losses of *Cetp* and *Eys*) have much higher values of % coding deletions and % coding substitutions compared to most other genes (in black). C) Histogram of the *MD* values for mouse genes along with the threshold values (dashed vertical lines) for conserved (✓, *MD* ≤ 5), undetermined (?), and likely eroded (†, *MD >* 15) genes. D) An example illustration of applying phylognetic filtering. We call hiconfErosion (×, e.g. in mouse and squirrel) if a gene is marked likely eroded (*MD >* 15) in more than one species in a clade that is flanked by at least one group in which the gene is conserved (✓). An undetermined gene in an individual species (?, e.g. in rat) is inferred as eroded if it resides in a clade with hiconfErosion. The likely eroded (†) gene in cow, however, is not considered a hiconfErosion since the finding is not supported by another adjacent species.

Importantly, this chain-based mapping approach succeeded for previously discovered gene losses, even accurately deciphering pseudogenized orthologs with extensive sequence erosion compared to the ancestral gene. For example, much of the exonic portions of *SERPINA10* are absent in the dog genome and do not yield a functional gene product [23]. As such, Ensembl does not offer a corresponding gene annotation in the dog genome assembly (canFam3), which would leave ambiguity about the existence and function this ortholog in dogs if not for experimental evidence. Our approach, however, precisely located the remnants of *SERPINA10* in the canFam3 assembly and allowed for assessing the semantic status (reading frame integrity and occurrence of substitution, insertion, and deletion) of the remaining chain-mapped exons (Figure 2C).

### Per-species identification of likely eroded orthologs using a Mahalanobis distance outlier model

Because functional inactivation will be accompanied by relaxation of purifying selection on all genomic regions encoding the trait, we used the extent of coding sequence erosion to declare the loss of gene function. In particular, to account for mutation rate heterogeneity across different branches of the mammalian tree, we restricted our results to orthologs that appear extremely eroded (using outlier detection) in sequence *relative* to all other mapped genes in the same given species. Unlike Sharma et al. [10], who used single-event features, such as in-frame stop codons and frameshift indels, we considered two different aggregate features of sequence erosion – i.e. the fraction of exonic nucleotides causing amino acid substitutions and the fraction out-right deleted with respect to the sequence of the human ortholog. This way, we avoided making spurious outlier calls caused by some mutations, such as stop codons near the end of a gene or multiple framshift indels that restore the original frame, which do not necessarily lead to loss of the ortholog’s function. Additionally, before declaring outliers, we masked out (i.e. did not consider) alignment gaps that could have resulted from sequencing gaps in the target species (see Methods).

For outlier detection, we used a Mahalanobis distance (*MD*) [24, 25] based model (see Methods) to mark a small fraction of the orthologs (typically 1-2% or 120-284 genes per species, median of 192; Figure 3A) as outliers for their extreme fraction both of non-synonymous substitutions and of deleted bases. We considered these outlier orthologs to be likely eroded in sequence and function in the species represented by the given genome assembly. For instance, by performing these methods on the 17,836 genes confidently mapped to the mm10 mouse assembly (Figure 3B), we distinguished previously discovered gene losses for *Cetp* [26] and *Eys* [27] (plus other novel predictions) as outliers (*MD >* 15), compared to uneroded, relatively sequence-conserved orthologs (*MD ≤* 5), including genes *Tp53* [28], *Notch1* [29], and *Sox11* [30] (Figure 3B,C) with indispensable function in mice.

### Phylogenetic filtering to identify high-confidence ortholog erosions in multiple related species

Two caveats must be considered for the per-species gene erosion predictions: Firstly, not all genes in the human reference set were necessarily present in functional form in the common ancestor shared by human and the a species under consideration. Sequences in other species that have some resemblance (such as paralogs) to a functional human gene, and that appear eroded, but were never present/functional in their common ancestor with human, could be confounded for gene erosion. Secondly, the erosion could be real but private to the single sequenced individual (i.e. not representative of the species at-large).

To address these caveats, we applied a number of strict phylogenetic filters to restrict the final candidate list to the highest-confidence predictions (Figure 3D). First, we required a final candidate to have an uneroded, sequence-conserved ortholog (*MD <* 5) in at least one other mammal in addition to human. With the assumption of parsimony, this first step ensures that the candidate ortholog was present, of functional importance, and under purifying selection in the common ancestor of human and any other extant lineages in which this ortholog is conserved in sequence. Second, even more importantly, we also required that each final candidate was considered likely eroded in *two or more* phylogenetically neighboring species (such as rat/mouse or manatee/elephant/cape elephant shrew) sharing a common ancestor with human in which the gene was declared conserved. Doing so minimized false positives from gene losses that are real but private to a sequenced individual, as well as those resulting from assembly, alignment, and mapping artifacts. We designated the final candidates as *<u>hi</u>gh-<u>conf</u>idence ortholog <u>erosions</u>*, or *hiconfErosions*.

In all, we identified 412 genes that were eroded in one or more multi-species groups, yielding 2,192 and 539 unique ortholog-species and ortholog-clade pairs, respectively (Figure 3A). Our hiconfErosion predictions include many previously known functional losses of *Cetp* [26], *Eys* [27], and *Tcn1* [31] in mouse, *ZP1* in bovine genomes [32], *CRYGB* in subterranean mammals [5], *GSDMA* in whales [10], and *RHBG* in fruit bats [10]. The number of hiconfErosions per species ranged from 5 in aardvark to 88 in brush-tailed rat and chinchilla (median of 41, Figure 4A).

**Figure 4.**
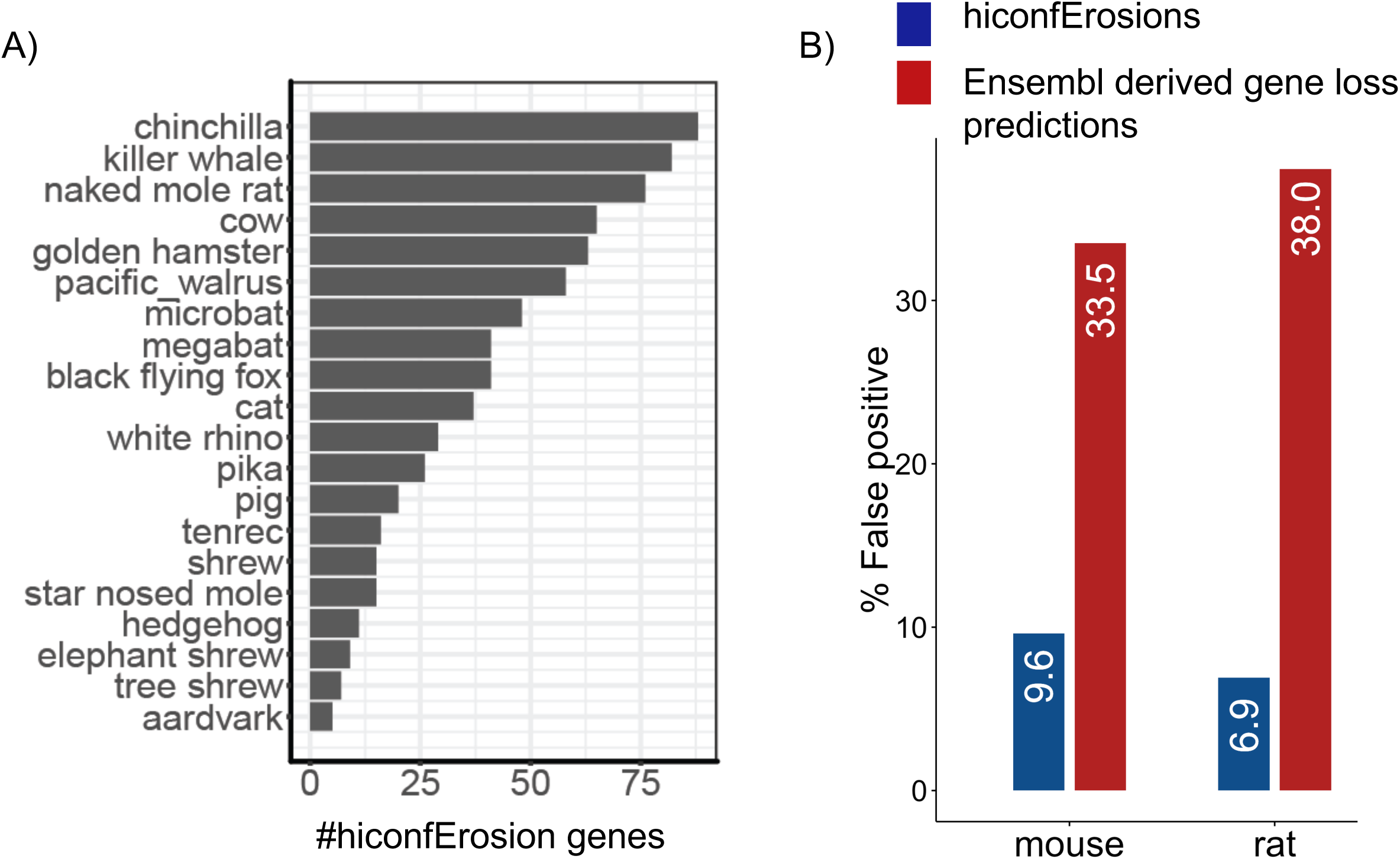
Distribution of hiconfErosions across species and estimation of False Positive Rate (FPR). A) Number of hiconfErosions predicted in different species. B) If hiconfErosions are directly inferred by applying the phylogenetic filtering (Figure 3D) to missing gene annotations in Ensembl (instead of eroded genes from Figure 3C), the estimated FPR (red bars) in mouse and rat is significantly higher than in our set of predicted hiconfErosions.

To estimate the false positive rate (FPR) for these results, we tabulated how many predicted hiconfErosions for mouse and rat were contradicted by proteomic or transcriptomic evidence from UniProt [18] (see Methods). We found 6/68 (8.82%) and 2/68 (2.94%) hiconfErosion predictions in mouse and rat conflicting with Uniprot evidence, respectively. We conservatively re-scaled the conflicts to account for potentially missing Uniprot entries but still found the FPR estimate to be less than 10% in both species (Figure 4B, see Methods). In contrast, if we applied our phylogenetic filtering directly to the orthologs missing Ensembl annotations for mouse and rat, the FPR for both species would be above 30% (Figure 4B), highlighting the importance of our orthologous chain mapping and outlier detection approach.

### Selection and variation in mammalian genes exhibiting multi-species ortholog erosion

Next, we asked how many hiconfErosions involve genes whose knockout mouse models do not survive beyond early development and, surprisingly, found ten such genes (Table 1, see Methods). We also found 27 hiconfErosions involve HGMD [33] genes implicated in severe human congenital disorders (Table 2, see Methods). Using the Residual Variation Intolerance Score (RVIS) [34], which is a gene-based measure of intolerance to common genetic variation (with minor allele frequencies greater than 5%), we found another 13 genes affected by hiconfErosions that have a relatively low RVIS score percentile (*<* 20%, Table 3, see Methods). We also found that our final hiconfErosion predictions (), containing 412 unique human orthologs, is depleted of genes causing lethality when inactivated in developing mice (p-value*<*2.2e-16; fold=8.43; Figure 5A), of known human HGMD disease-causing genes (p-value*<*1.3e-11, fold=3.17; Figure 5B), and of those intolerant to functional variation in humans (p-value*<*1e-6, Figure 5C).

**Figure 5.**
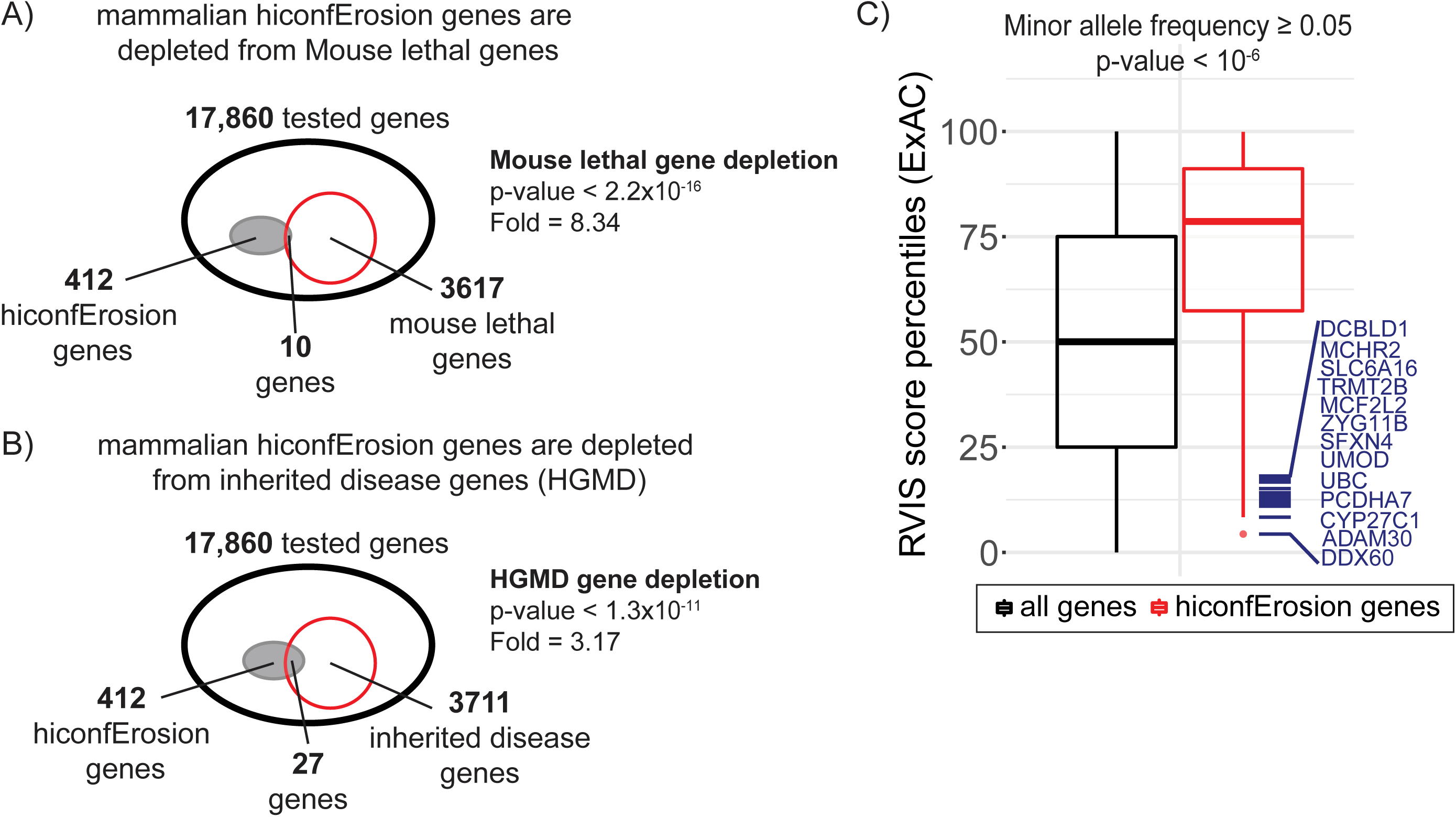
Selection and variation in mammalian genes affected by hiconfErosion. A) hiconfErosion genes are depleted from genes whose inactivation in mouse models leads to lethal consequences in early development. B) hiconfErosion genes are depleted from genes with known Mendelian disease mutations (HGMD genes). C) We compare the genome-wide distribution of Residual Variation Intolerance Score (RVIS) percentiles (derived from ExAC) for common variation at MAF of 0.05 to the RVIS percentiles of genes affected by hiconfErosions. By and large, hiconfErosion genes are more tolerant of common functional variation in humans, suggesting that selection over coding genes is similar across mammalians. Several genes at the lower tail of the RVIS scores are highlighted (see Table 3 for details).

**Table 1.**
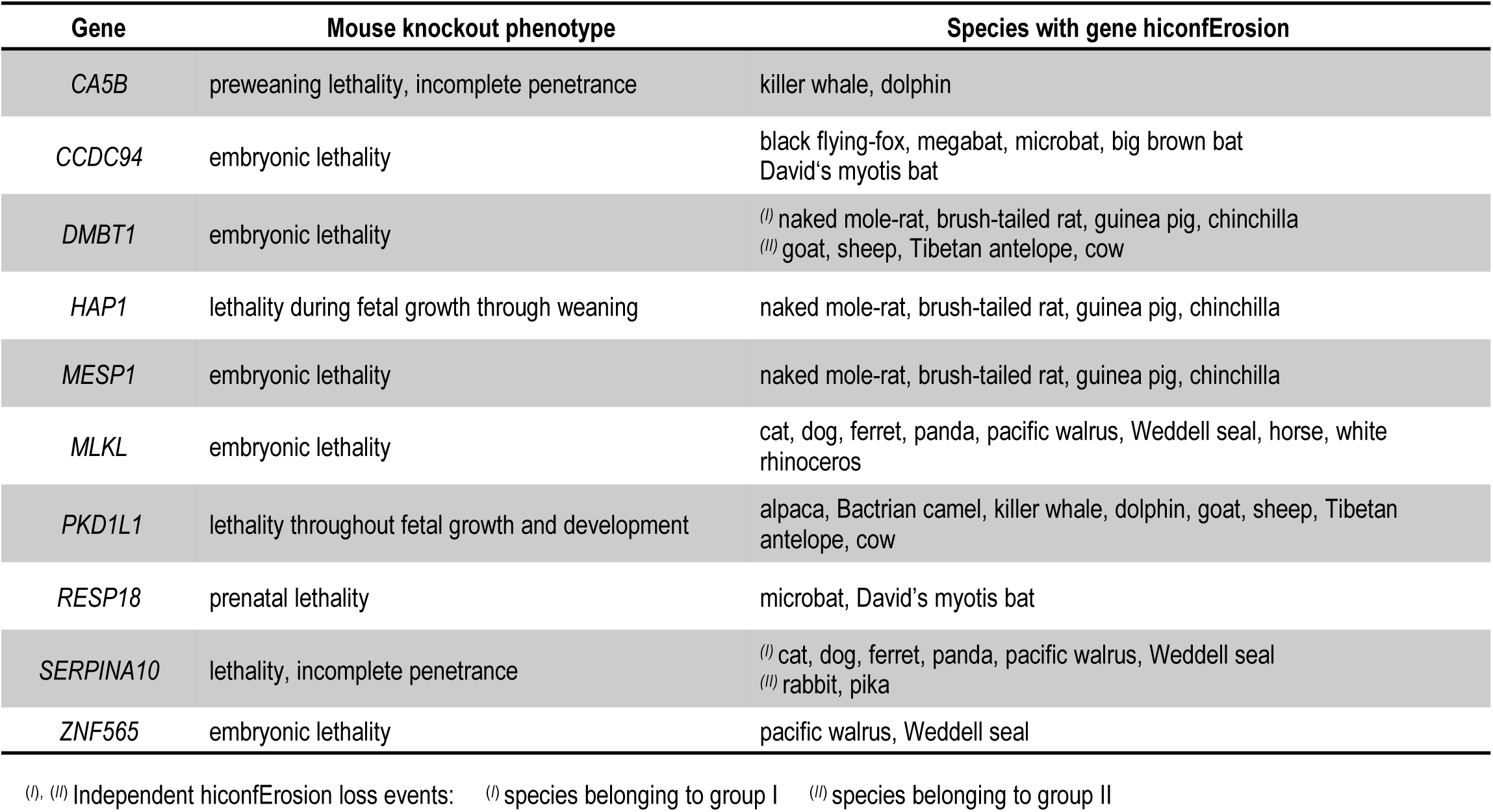
Genes implicated in mammalian hiconfErosions that result in lethality when inactivated in mouse models.

**Table 2.** Genes implicated in mammalian hiconfErosions that harbor monogenic disease variants in human patients (disease mutations in HGMD [33]).

| Gene | Associated human monogenic disease | Species with gene hiconfErosion |
| --- | --- | --- |
| *A2ML1* | Noonan syndrome<br>Otitis media (susceptibility to) | naked mole-rat, brush-tailed rat, guinea pig, chinchilla |
| *CETP* | Higher/Lower HDL cholesterol level<br>Hyperalphalipoproteinaemia<br>Cholesterol ester transfer protein deficiency | <sup>(I)</sup> Bactrian camel, alpaca<br><sup>(II)</sup> goat, sheep, Tibetan antelope, cow |
| *CRYBB3* | Cataract | microbat, big brown bat, David's myotis bat |
| *CRYGB* | Congenital cataract | cape golden mole, tenrec |
| *EYS* | Retinitis pigmentosa<br>Retinal dystrophy<br>Peripheral dystrophy<br>Usher syndrome | mouse, rat, Chinese hamster, golden hamster, prairie vole |
| *FCN2* | FCN2 deficiency | goat, sheep, Tibetan antelope, cow |
| *GCKR* | Elevated HDL-cholesterol<br>Hypertriglyceridaemia | microbat, big brown bat, David's myotis bat |
| *HMGCS2* | Mitochondrial HMG-CoA synthase deficiency | black flying-fox, megabat |
| *HSD3B2* | 3 beta-hydroxysteroid dehydrogenase deficiency<br>Adrenal hyperplasia<br>Hypospadias, non-syndromic<br>Idiopathic hypospadias<br>Pseudohermaphroditism | goat, sheep, Tibetan antelope, cow |
| *KLKB1* | Prekallikrein deficiency | dolphin, killer whale |
| *KRT3* | Corneal dystrophy, Meesmann | mouse, rat |
| *MESP1* | Tetralogy of Fallot<br>Ventricular septal defect | naked mole-rat, brush-tailed rat, guinea pig, chinchilla |
| *MMP21* | Heterotaxy | pig, dolphin, killer whale, Bactrian camel, alpaca, goat, sheep, Tibetan antelope, cow |
| *MYD88* | MYD88 deficiency | brush-tailed rat, chinchilla |
| *OPTC* | Glaucoma, primary open angle | cape golden mole, tenrec, microbat, big brown bat, David's myotis bat |
| *PCSK9* | Hypercholesterolaemia<br>Low/High LDL cholesterol | <sup>(I)</sup> cat, dog, ferret, panda, pacific walrus, Weddell seal<br><sup>(II)</sup> goat, sheep, Tibetan antelope, cow |
| *PLA2G2A* | Colorectal cancer | dolphin, killer whale |
| *RIPPLY1* | Intellectual disability | black flying-fox, megabat |
| *SFXN4* | Colorectal cancer<br>Mitochondriopathy and macrocytic anaemia | <sup>(I)</sup> Chinese hamster, golden hamster, prairie vole<br><sup>(II)</sup> naked mole-rat, brush-tailed rat, guinea pig, chinchilla |
| *STAP1* | Hypercholesterolaemia | dolphin, killer whale |
| *TCN1* | Transcobalamin I deficiency | mouse, rat |
| *TNFRSF6B* | Systemic lupus erythematosus | mouse, rat, Chinese hamster, golden hamster, prairie vole |
| *UMOD* | Chronic kidney disease<br>Congenital anomalies of the kidney and urinary tract<br>FJHN/MCKD syndrome<br>Glomerulocystic kidney disease<br>Uromodulin-associated kidney disease<br>Hyperuricaemic nephropathy | microbat, big brown bat, David's myotis bat, black flying-fox, megabat |
| *XAF1* | Multiple sessile serrated adenoma | cat, dog, ferret, panda, pacific walrus, Weddell seal |
| *ZNF543* | IgA nephropathy | cat, dog, ferret, panda, pacific walrus, Weddell seal |
| *ZNF589* | Intellectual disability | ferret, panda, pacific walrus, Weddell seal |
| *ZP1* | Infertility | pig, dolphin, killer whale, Bactrian camel, alpaca, goat, sheep, Tibetan antelope, cow |

(I), (II) Independent hiconfErosion loss events: (I) species belonging to group I (II) species belonging to group II

**Table 3.** Genes implicated in mammalian hiconfErosions that have low RVIS (Residual Variation Intolerance Score) score percentile. The RVIS scores were inferred based on Exome Aggregation Consortium variants with Minor Allele Frequency of 0.05%.

| Gene | RVIS Score Percentile | Species with gene hiconfErosion |
| --- | --- | --- |
| *DDX60* | 4.36% | Tibetan antelope, Sheep, Domestic goat, Cow |
| *ADAM30* | 8.36% | Guinea Pig, Brush-tailed rat, Naked mole-rat, Chinchilla |
| *CYP27C1* | 10.99% | David's myotis (bat), Microbat, Big brown bat |
| *PCDHA7* | 11.82% | Panda, Weddell seal, Ferret, Pacific walrus |
| *UBC* | 12.52% | <sup>(I)</sup> Hedgehog, Star-nosed mole, Shrew<br><sup>(II)</sup> Naked mole-rat, Guinea Pig, Chinchilla, Brush-tailed rat<br><sup>(III)</sup> Ferret, Panda, Pacific walrus, Weddell seal<br><sup>(IV)</sup> Alpaca, Bactrian camel |
| *UMOD* | 13.17% | Black flying-fox, Megabat, David's myotis (bat), Microbat, Big brown bat |
| *SFXN4* | 13.78% | <sup>(I)</sup> Chinese Hamster, Golden hamster, Prairie vole<br><sup>(II)</sup> Naked mole-rat, Guinea Pig, Chinchilla, Brush-tailed rat |
| *ZYG11B* | 14.17% | Rhesus macaque, Crab-eating macaque |
| *MCF2L2* | 15.17% | Golden hamster, Mouse, Rat, Lesser Egyptian jerboa, Prairie vole, Chinese Hamster |
| *TRMT2B* | 16.72% | Guinea Pig, Chinchilla, Naked mole-rat, Brush-tailed rat |
| *SLC6A16* | 17.45% | Black flying-fox, Megabat |
| *DCBLD1* | 18.09% | Lesser Egyptian jerboa, Prairie vole, Golden hamster, Chinese Hamster, Mouse, Rat |
| *MCHR2* | 18.15% | Naked mole-rat, Guinea Pig, Chinchilla, Brush-tailed rat |

<sup>(I)</sup>, <sup>(II)</sup>, <sup>(III)</sup>, <sup>(IV)</sup> Independent hiconfErosion loss events: <sup>(I)</sup> species belonging to group I <sup>(II)</sup> species belonging to group II <sup>(III)</sup> species belonging to group III <sup>(IV)</sup> species belonging to group IV

### Erosion of *GCKR* in microbats may facilitate their high-fat insectivorous diet

Microbats and megabats, the two primary suborders of bats, have vastly different diets. Microbats have extremely fat-rich, insect-based diets [35], while megabats have fruit-based diets depleted of fat [36]. We found that *GCKR* (Glucokinase regulatory protein; OMIM: 600842), variants of which are associated with elevated high-density lipoprotein (HDL) cholestrol in human [37], is specifically eroded in microbats (see Figure 6 and Table 2). This potentially provides a genotypic basis for an observation by Widmaier et al. [38] who found that HDL-cholestrol levels in a lactating microbat species (*Tadarida brasiliensis mexicana*) were 10-fold higher in comparison to three species of megabats. Specifically, they proposed that these extraordinarily high HDL levels, possibly facilitated by *GCKR* erosion, could explain why microbats bats are protected from atherosclerosis and other common diseases associated with high-fat intake.

**Figure 6.**
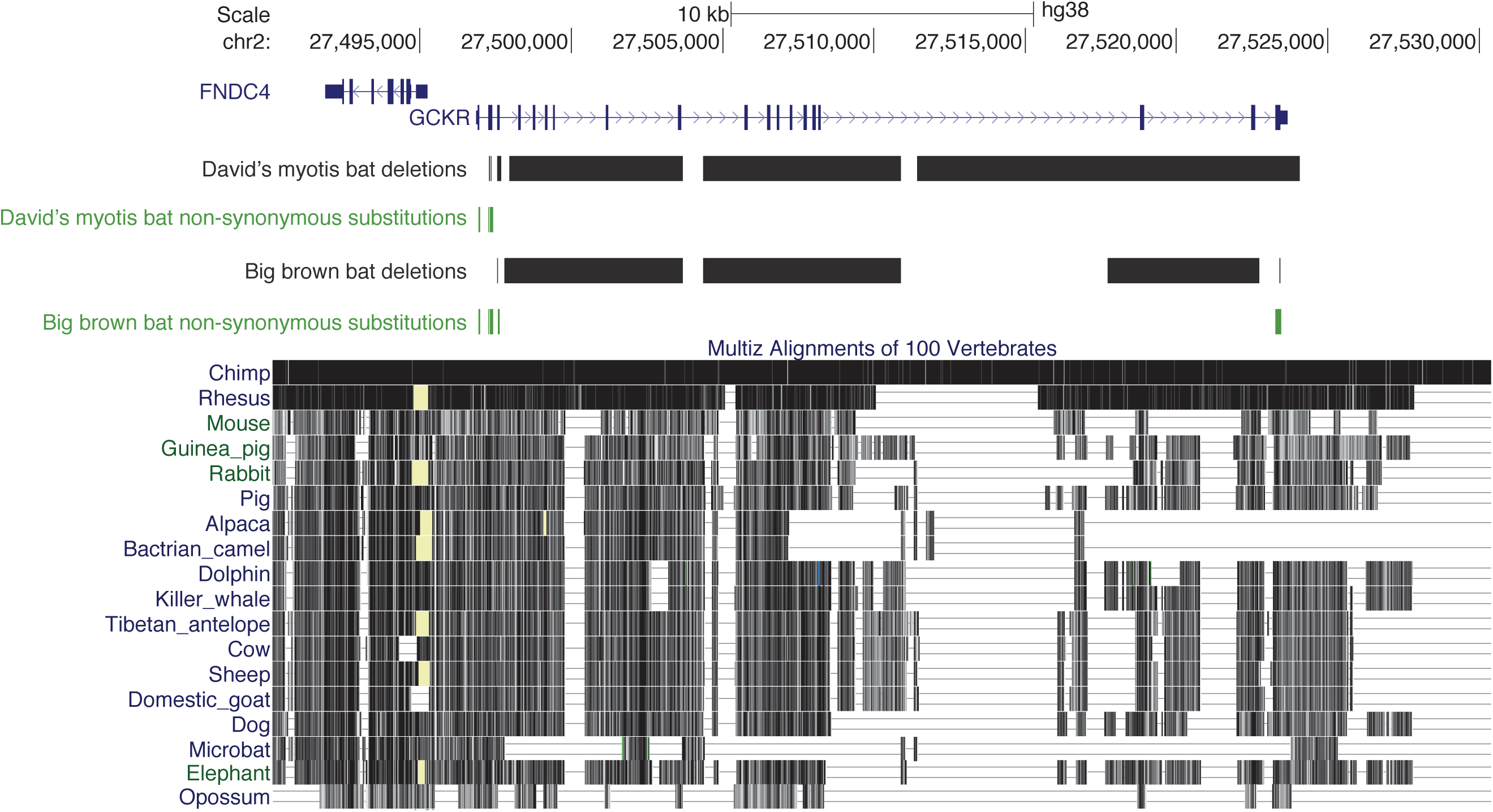
*GCKR* hiconfErosion in microbats. A genome browser view of *GCKR* (Glucokinase regulatory protein) on the human genome (GRCh38) and a mammalian multiple alignment highlight coding lesions in microbat. Positions of deleted genomic fragments (black) and non-synonymous substitutions (green) are highlighted for David’s myotis and big brown bat (both of *microchiroptera*).

### Erosion of *PKD1L1* and *MMP21* evolved in artiodactyla (even-toed ungulates) but causes prenatal lethality in mice and congenital heart defects in humans

*Pkd1l1* (polycystin kidney disease protein 1-like 1) null mutations in mice result in a high rate of embryonic lethality [39] (Table 1) characterized by misregulated nodal signaling [40] and anomalous Left-Right (L-R) symmetry patterning [41, 42] (Figure 7A). Indeed, humans with homozygous loss of function in *PKD1L1* (OMIM: 609721) present with laterality defects ranging from *Situs Inversus Totalis* to heterotaxy and with congenital heart disorders, all of which can be deleterious early in life [43]. As such, it was remarkable to find *PKD1L1* hiconfErosion, with deletion of as much as 98% of coding bases in some species, in a clade of 9 even-toed ungulates (including cetaceans, see Figure 7B). Congruent with this finding, an Ensembl annotation for *PKD1L1* does not exist for the cow, sheep, dolphin, alpaca and pig genomes, and we detected no orthology between human *PKD1L1* to any other gene in these genomes. In contrast, we found high sequence homology for the human protein sequence of *PKD1L1* in the genomes of outgroup lineages such as horse and elephant, suggesting that the gene is intact in those species, and lost in artiodactyla (even-toed ungulates).

**Figure 7.**
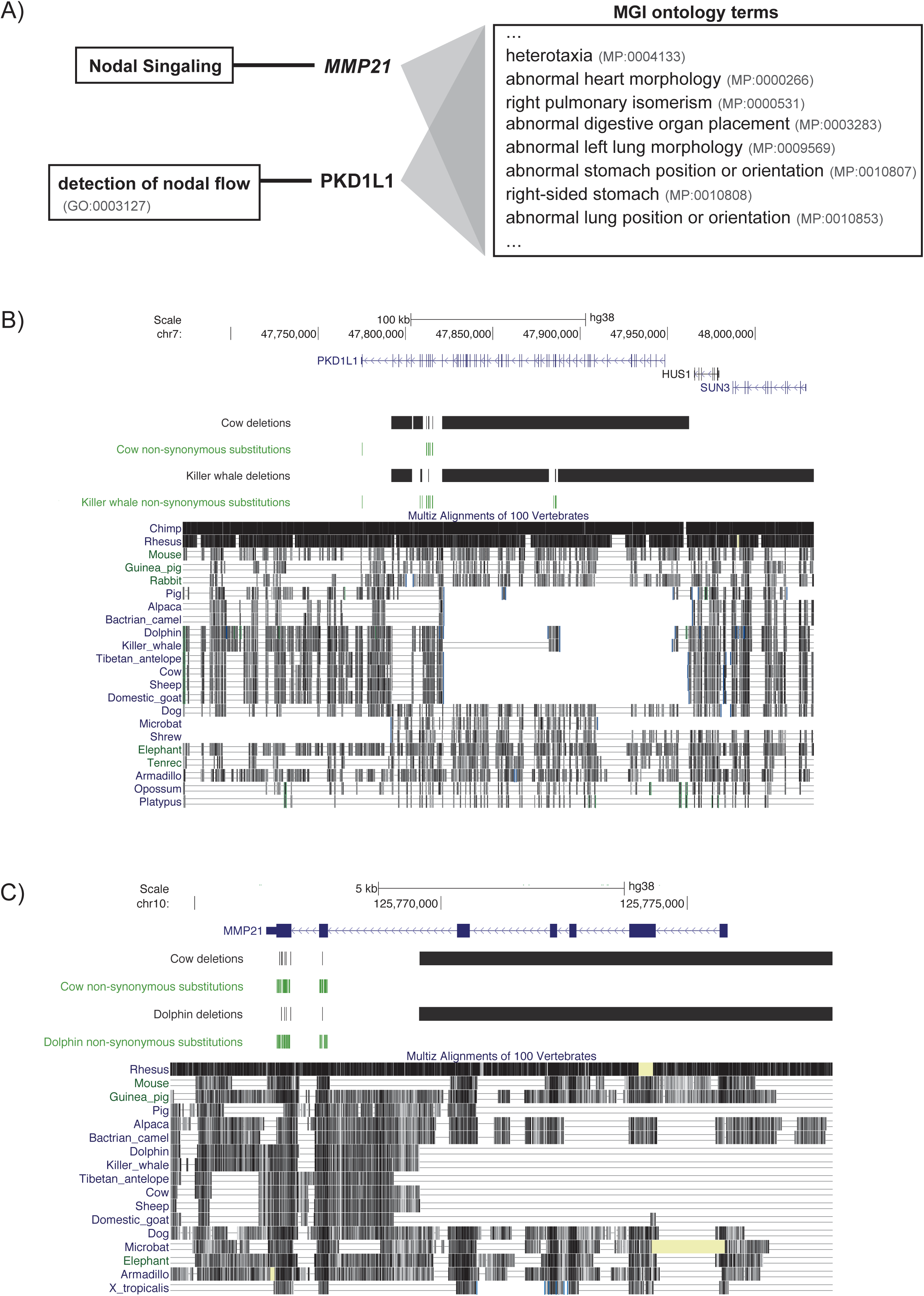
*PKD1L1* and *MMP21* hiconfErosions in even-toed ungulates. A) *PKD1L1* (Polycyctic kidney disease protein 1-like 1) and *MMP21* (Matrix Metalloproteinase 21) are both related to the Nodal signaling pathway, and null mutations in either gene results in severe laterality disorders with congenital heart defects. B) A genome browser view of *PKD1L1* (on the human genome, GRCh38) and a mammalian multiple alignment highlight a clade-specific lesion of the gene in 9 even-toed ungulate species. Positions of deleted fragments (black) and non-synonymous substitutions (green) are highlighted for cow and killer whale. C) *MMP21* has undergone hiconfErosion in the same ancestor. Positions of deleted fragments (black) and non-synonymous substitutions (green) are highlighted for cow and dolphin.

Interestingly, we also discovered a related hiconfErosion in the same group of even-toed ungulates (including cetaceans) affecting the gene *MMP21* (Matrix Metalloproteinase 21; OMIM: 608416; Figure 7C) that encodes components of Nodal signaling [43] and is also implicated in heterotaxy [44] and in congenital heart disorders [45, 46] (Figure 7A). Interestingly, Double Outlet Right Ventricle – a cardiac disorder associated with mutation in mouse *Mmp21* [47] and with mutation in human *PKD1L1* [43] – has significantly higher incidence in cattle with congenital heart disorder relative to human and other animals with this condition [48] (presumably because hiconfErosion of *both PKD1L1* and *MMP21* naturally occurs in artiodactyla but not in other mammals).

### Erosion of endocrine regulator *GPRC6A* evolved in odontoceti (toothed whales) but causes testicular insufficiency in humans and mice

*GPRC6A* (OMIM:613572) is a G protein-coupled receptor that plays an important role in metabolic and endocrine regulation [49]. Highly expressed in Leydig cells of the testis (in addition to many other tissues), *GPRC6A* is activated by testosterone, osteocalcin, basic amino acids, and various cations [50, 51]. In mouse models, *Gprc6a* ablation results in dramatic decrease of testosterone levels, smaller testis size, and reduced sperm count [52]. Similarly in humans, loss-of-function mutations in *GPRC6A* are associated with testicular insufficiency (subfertility, altered sperm parameters, low circulating testosterone levels, and high circulating luteinizing hormone levels). A number of studies also found that *GPRC6A* is associated with prostate cancer progression and could serve as a potential drug target for this malignancy [51, 53].

*GPRC6A* is a highly conserved gene across mammals and even has orthologs in species as distant as zebrafish (Figure 8), suggesting it may have arisen early in vertebrate evolution. It is highly remarkable, then, that we observed hiconfErosion of *GPRC6A* in dolphin and killer whale – where over 50% of coding bases are deleted in both species (including the entirety of exons 5 and 6), and the remaining regions contain several frameshift indels and nonsense mutations (Figure 8). The gene is otherwise well-conserved in bovidae (domestic goat, sheep, Tibetan antelope, and cow), suggesting it was functional in the cetartiodactyla ancestor (superorder of odontoceti and bovidae) but was inactivated and then allowed to erode in the evolutionary past of toothed whales. *GPRC6A* erosion in toothed whales is especially striking when considering that toothed whales exhibit 7-25 times greater testis-to-body mass ratio relative to other mammals [54].

**Figure 8.**
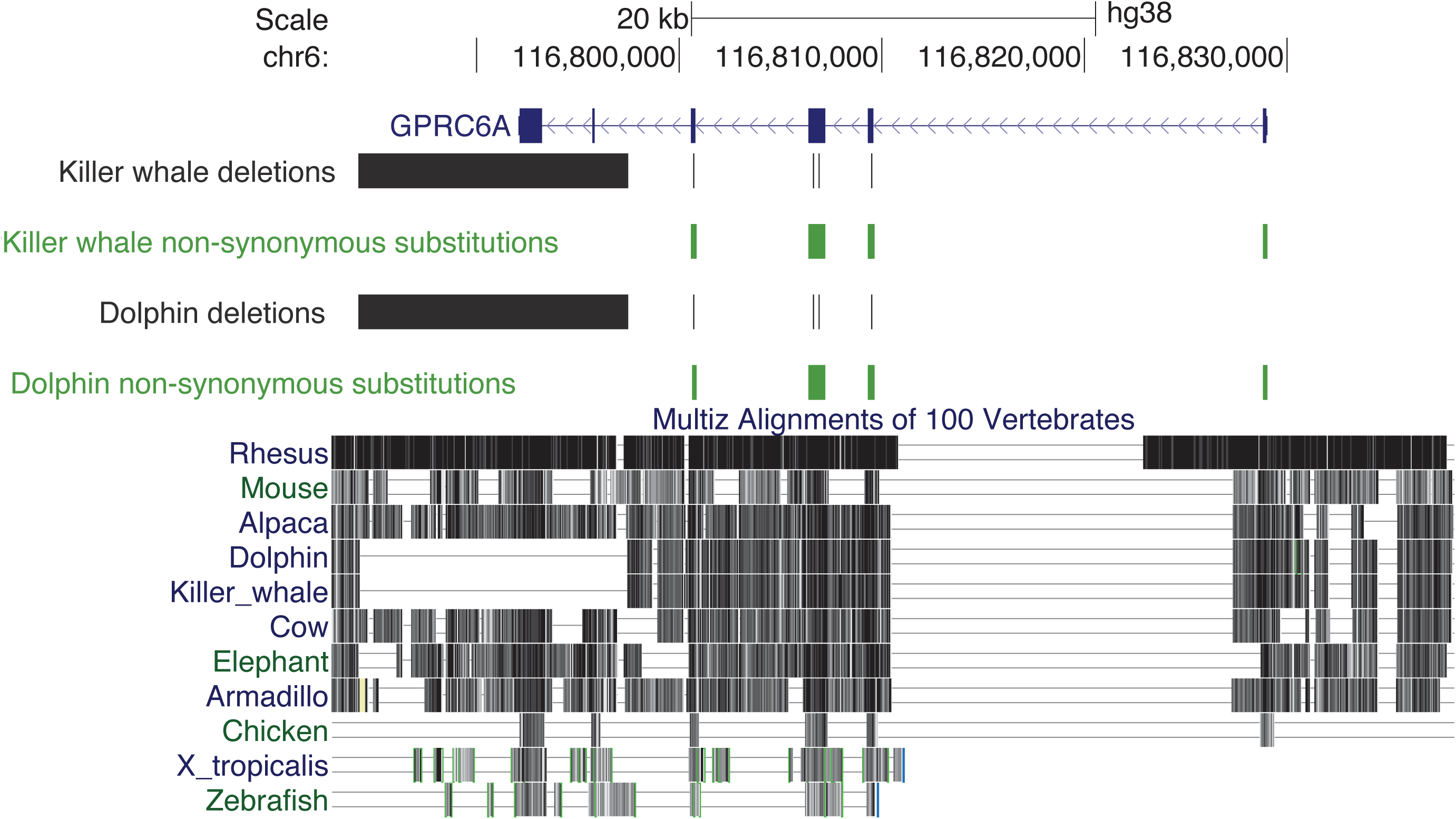
*GPRC6A* hiconfErosion in toothed whales. A genome browser view of *GPRC6A* (on the human genome, GRCh38) and a mammalian multiple alignment highlight coding lesions in dolphin and killer whale. Positions of deleted fragments (black) and non-synonymous substitutions (green) are shown for both species. *GPRC6A* is a key hormonal and metabolism regulation protein, and the knockout of *GPRC6A* in male mice results in over 6-fold decrease of testosterone levels and reduced size and weight of the testis [52].

## Discussion

In this work, we devised a novel approach to automate the survey of genomes for instances of coding gene sequence erosion. Knowledge of gene losses has far-reaching applications in studying the morphological and physiological adaptations of different species [10], in discovering alternative molecular pathways for critical genes [2], in medical genetics [13], and even in animal conservation programs [12]. We designed our approach for low false positive rates in order to be practically useful for the experimentalists tackling these research efforts. In particular, our conservative approach addresses a number of challenges that hindered previous automated methods.

Pre-existing genome annotation methods [17] were designed to identify intact genes and were not tuned to distinguish sequence erosion due to a bonafide gene loss from confounds due to alignment, assembly, and other technical artifacts. In contrast, our mapping procedure using whole-genome pairwise alignments is synteny-aware, is robust to sequencing gaps and other artifacts, and produces strictly one-to-one ortholog assignments that are not confounded by gene duplication events (paralogs) across species. For each genome, we used the mapped gene annotations to scan for signatures of gene erosion (large deletions and non-synonymous substitutions) [10] while also carefully accounting for sequencing gaps and other assembly artifacts. To enrich for high-confidence gene erosion predictions, we used a Mahalanobis distance-based metric to refine the candidates to just those orthologs exhibiting extreme amounts of deletion and non-synonymous substitution relative to other genes in that species. We designated these orthologs as being likely eroded in sequence and function in the species represented by each examined assembly. To even further minimize erroneous predictions, we distilled out just those candidates that are considered likely eroded in at least two closely related species but sequence-conserved in at least one other outgroup besides human. Despite the phylogenetic filter reducing sensitivity for detecting gene losses, we discovered 2,192 orthologs (hiconfErosions) across 52 species that are severely eroded. Many of these hiconfErosions are novel candidates with evolutionary and disease significance (Table 1 and Table 2). Evaluated against UniProt expression and proteomic observations in mouse and rat (see Methods), this final candidate set has an estimated false prediction rate of less than 10%.

While a majority of the hiconfErosions were predicted to have benign consequences, dozens involved loci associated with lethal developmental outcomes in mouse knockout models (Table 1), human disease (Table 2), and/or intolerance to variation in human populations (Table 3). The evolutionary mechanisms contributing to these functionally significant gene losses were likely diverse. Some may have resulted from regressive evolution (i.e., degeneration of formerly useful anatomical and physiological functions), such as the degraded *OPTC* vision gene [55] in nocturnal microbats or the *CRYGB* erosion in subterranean cape golden moles. Other losses may actually have contributed to species adaptation, such as erosion of the epithelium gene *GSDMA* [56, 57] in cetaceans which have skin adaptations to life in the water [10].

Species with hiconfErosion that tolerate functional changes known to cause disease in humans provide intriguing potential for medical genetics [13]. Some of these “knockout experiments of nature” may tolerate and even derive selective advantage from gene loss(es) because of factors specific to their habitat niche. Others, however, may have evolved compensatory suppressor mutations to genetically negate otherwise deleterious consequences. It is these latter cases that could be harnessed for treating human congenital disorders [13]. For instance, despite the erosion of *GPRC6A* in toothed whales, these magnificent creatures have relatively massive testes for their size – the categorically opposite phenotype of when this gene is inactivated in mice and humans. We posit that further examination of odontoceti could reveal complementary testis development pathways that might inform therapy for male hypogonadism. More broadly, extension of this gene loss survey to additional or newly sequenced species across the metazoan tree of life will likely reveal more such evolutionary curiosities with great clinical potential.

To that end, we emphasize the ease with which genomes can be added to analyses using our approach. Each additional genome would independently be aligned to the reference assembly – without the high computational cost of realigning all genomes, as is necessary for methods requiring multiple-sequence alignments. Thus, our approach is easily scalable for the ever-increasing number of new species being sequenced [15, 58].

## Methods

### Genome assemblies, reference genome and protein-coding gene annotations

We used genome assemblies of 59 mammalian species (listed in Supplementary Table 1) in this study, with human genome assembly GRCh38/hg38 as reference. From Ensembl biomart release 86, we downloaded lists of gene identifiers and counted how many unique ‘protein coding’ gene identifiers are found. Of the 59 mammalian species, 34 species had gene annotation in Ensembl. For our reference gene set, we started with 22,072 unique human protein-coding genes annotated in Ensembl and excluded gene annotations from un-placed and un-localized scaffolds, leaving 19,729 protein-coding genes. We then obtained coordinates for segmental duplications (genomic fragments larger than 1kb with more than 90% sequence identity to other genomic fragments) from the UCSC genome browser and excluded from our analysis genes with 10% or more overlap with any segmentally duplicated fragment. We also restricted the reference gene set to genes having “complete” ENSEMBL transcripts. These filters generated our reference gene set of 17,860 unique human protein-coding genes for mapping across mammalian genomes. We collected assembly N50 measures for each genome from NCBI: https://www.ncbi.nlm.nih.gov/assembly.

### Mapping orthologs across mammalian genomes

We used Jim Kent’s BLASTZ-based, whole-genome pairwise alignment chains [22] to map – when possible with high confidence – each reference gene to a single orthologous position in each of the 58 query species (Figure 2A). First, we assigned every base in the canonical transcript of the reference gene to the highest-scoring chain (in terms of UCSC chain alignment scores) that contains the base in its alignment (Figure 2B1-B2). The chain to which most number of bases get assigned was picked as the *best chain* (*C_b_*) for that gene (Figure 2B3). Since chains are preferentially ordered based on the alignment score before assigning bases (Figure 2B1), the best chain was also the chain with highest synteny for those aligning bases, and was treated as the most likely ortholog for that gene. Next, this procedure was repeated after removing the *best chain* (*C_b_*) from consideration to pick another chain as the *second-best chain* (*C_sb_*) for the same gene (Figure 2B4). This chain *C_sb_* was treated as containing a likely paralog of the gene. To ensure that the orthologous chain was easily differentiable from paralogs, we required the *second-best chain* to have an alignment score at least 20 times lower than that of the best chain. To also ensure high synteny, we required the number of bases in the aligning blocks of the best chain be at least 20 times greater than the number of bases in the gene itself, i.e. *gene-in-synteny ≥* 20, where *gene-in-synteny* = length of *C_b_* / length of gene. We also required unique mapping of coordinates between reference and query genomes, such that if two or more reference genes were mapped to the same location, all overlapping mappings were discarded.

In each species, for every gene to which an orthologous chain was assigned, we used the alignments derived from the chain to determine the orthologous amino acid sequence for every transcript of that gene. Because exon boundaries sometimes shift for evolutionary or alignment reasons, we excluded the first and last two amino acid positions in each exon for all downstream analyses [59]. Gaps in chain alignments found in regions containing an assembly gap in the query species were masked out to avoid confusing them with true deletions in the query genome.

### Outlier detection model

To identify the genes orthologs that have a surprisingly high number of non-synonymous coding substitutions and deletions in each of the 58 query species, we used an outlier detection approach. We first used the derived amino acid sequence of each transcript for every gene in a query species, as described previously, to compute a feature vector 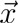 = (*x*_1_*, x*_2_), where *x*_1_ is the fraction of the transcript gene deleted in the query species and *x*_2_ is the fraction of non-synonymous coding substitutions in the remaining intact portion of the gene transcript compared to the orthologous gene transcript sequence in the reference species (human).

We then used 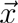 across all genes in the query species to compute the mean vector, 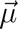, and a covariance matrix, *S*. Mahalanobis distance (*MD*) [24, 25] measures how “far” a given gene with feature vector, 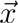, is from the distribution of all genes in the query species, as follows:

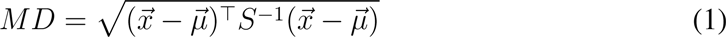

Since most genes exhibit relatively few non-synonymous substitutions and deletions (as placental mammals are relatively closely related, see Figure 3B), a very large value of *MD* implies high extensive erosion of the gene. Conversely, a very small value of *MD* implies that the gene is well- conserved. Since the coding sequences are expected to diverge more with increasing phylogenetic distance between reference and query species, we computed a different distribution (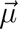 and *S*) for each query species. If all transcripts of an ortholog had *MD* greater than an absolute threshold (*MD >* 15), we marked the gene an outlier (✓) since it is likely sequence-eroded and nonfunctional in the species represented by the query genome. If all transcripts of an ortholog exhibited *MD ≤* 5, we regarded the gene sequence-conserved and functional (✓) in the species represented by the query genome. All other orthologs that were not categorized into either of the former two groups were marked with undetermined (?).

### Phylogenetic filtering

We used the phylogenetic tree of the 59 mammalian species (including human; Supplementary Figure 1) to identify which of the outlier orthologs would be considered to have undergone hiconfErosion (Figure 3D illustrates this procedure). For each gene, we recorded which species exhibit a sequence-conserved ortholog (based on the results of the outlier detection model described above; also see Figure 3B). Since the gene under consideration is already annotated to be functionally present in hg38 by Ensembl, we inferred by parsimony that the last common ancestor of human and of all species containing the conserved gene likely had a functional copy of this gene with a sequence similar to that of its extant functional orthologs. Finally, we search all subtrees starting from the earliest ancestral species marked in this fashion with gene conservation to identify specific groups where hiconfErosion has occurred. A group in which hiconfErosion has occurred is one in which there are at least two species with the gene marked as likely eroded (all transcripts with *MD >* 15) and no leaf-node species with the gene marked as sequence-conserved (*MD ≤* 5).

### Curation of Uniprot data and details of FPR (False Positive Rate) estimation in mouse and rat

To measure the rate of false positives in hiconfErosion predictions for mouse and rat, we obtained experimentally curated coding gene models downloaded as protein sequences with ‘protein level’ or ‘transcript level’ evidence from Uniprot [18] (release-2017 01; ftp://ftp.uniprot.org/pub/databases/uniprot/current_release/knowledgebase/complete).

We found 16,385 and 7,591 gene models in mouse and rat, respectively. We then mapped those amino acid sequences to their respective genomes (mouse sequences to mm10; rat sequences to rn6) using protein BLAT [60] with default settings (minimum mapping score 30; minimum sequence identity 25; maximum intron size 750,000). After mapping these protein sequences, we computed what proportion of orthologs predicted to be affected by hiconfErosion is contradicted by empirical evidence. Specifically, for a given species and its genome, any predicted hiconfErosion-affected orthologs that deviate by more than 20% in sequence length, share less than 90% identity, or disagree with the locus of the relevant Uniprot sequence were considered false predictions. Lastly, since we had 17,860 tested proteins in the reference gene set, we conservatively scaled up the number of conflicts in mouse and rat by a factor of 1.09 (=17,860/16,385) and 2.35 (=17,860/7,591), respectively, to estimate our False Positive Rate (FPR) while accounting for potentially missing Uniprot entries.

### Curation of lethal loss of function genes in mouse models

We used gene function annotations from the MGI Phenotype Ontology (Mouse Genome Informatics, http://www.informatics.jax.org/) to derive a subset of genes whose disruption leads to severe developmental perturbations and lethal consequences. The MGI Phenotype Ontology catalogs spontaneous, induced, and genetically-engineered mouse mutations and their associated phenotypes. Ontology data (containing 8,949 phenotypic terms; v6.04) were lifted over from mouse to human, resulting in 609,253 (canonicalized) gene-phenotype associations. We selected all 50 unique phenotypic terms with the string “lethal” in their description (e.g., MP:0008762 *embryonic lethality*; MP:0008569 *lethality at weaning*; MP:0006206 *embryonic lethality between somite formation and embryo turning*), identifying a total of 3,617 genes associated with causing early lethality when disrupted.

### Analysis of hiconfErosion genes implicated in HGMD

We obtained an access to a professional version (2016.2) of the Human Gene Mutation Database [33] containing 165,939 entries of known disease-causing gene mutations (referenced to GRCh38 assembly) and their respective phenotypes. We then obtained the list of 4,014 monogenic disease genes (i.e. genes with at least one “DM” mutation) from this database, of which 3,711 genes were also present in our reference gene set.

### Analysis of genic variation intolerability with RVIS score

Residual Variation Intolerance Score (RVIS) is a unified genome-wide metric that quantifies the extent to which genes tolerate coding mutation load (i.e., the higher the score, the more a gene can tolerate common genetic variation without negative consequences). We downloaded the precomputed percentiles of RVIS scores (inferred based on Exome Aggregation Consortium variants with Minor Allele Frequency of 0.05%) available for 17,276 genes (http://genic-intolerance.org/data/RVIS_Unpublished_ExAC_May2015.txt). We recorded the hiconfErosion genes with low (*<* 20%) RVIS score percentiles, which are relatively intolerant to functional variation in humans.

### Statistical significance of depletion

To estimate statistical significance of depletion [61] for the hiconfErosion group of genes relative to categories such as *mouse lethal genes* or *HGMD disease genes*, we used the Fisher exact test to compute a one-sided p-value (using R):

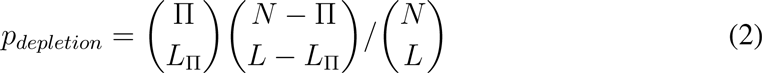

In the above equation, *N* and *L* denote the total number of genes and the hiconfErosion subset of genes in the analysis, respectively. Similarly, Π and *L*_Π_ denotes the subset of *N* and *L* associated with a gene category such as HGMD disease genes or mouse lethal genes.

We also compute a ‘fold change’ for depletion as 1*/Fold_enrichment_*:

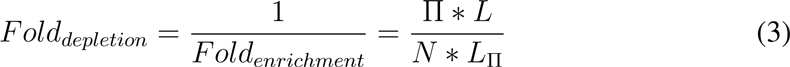

We calculate the two-tailed p-value using R to test how different were the mean RVIS scores of the 412 genes implicated in hiconfErosions compared to the reference gene set.

## Acknowledgements

We thank the members of the Bejerano Laboratory, particularly M. J. Berger, K. Jagadeesh, J. Birgmeier, B. Yoo and others for technical advice and helpful discussions. We thank Hiram Clawson and the UCSC Genome Browser team for providing us the mammalian alignment chains and multiple sequence alignments and technical advice. We thank D. Cooper and P. Stenson for Human Gene Mutation Database (HGMD) access.

## Competing Interests

The authors declare that they have no competing financial interests.

---

**Supplementary Table 1.**
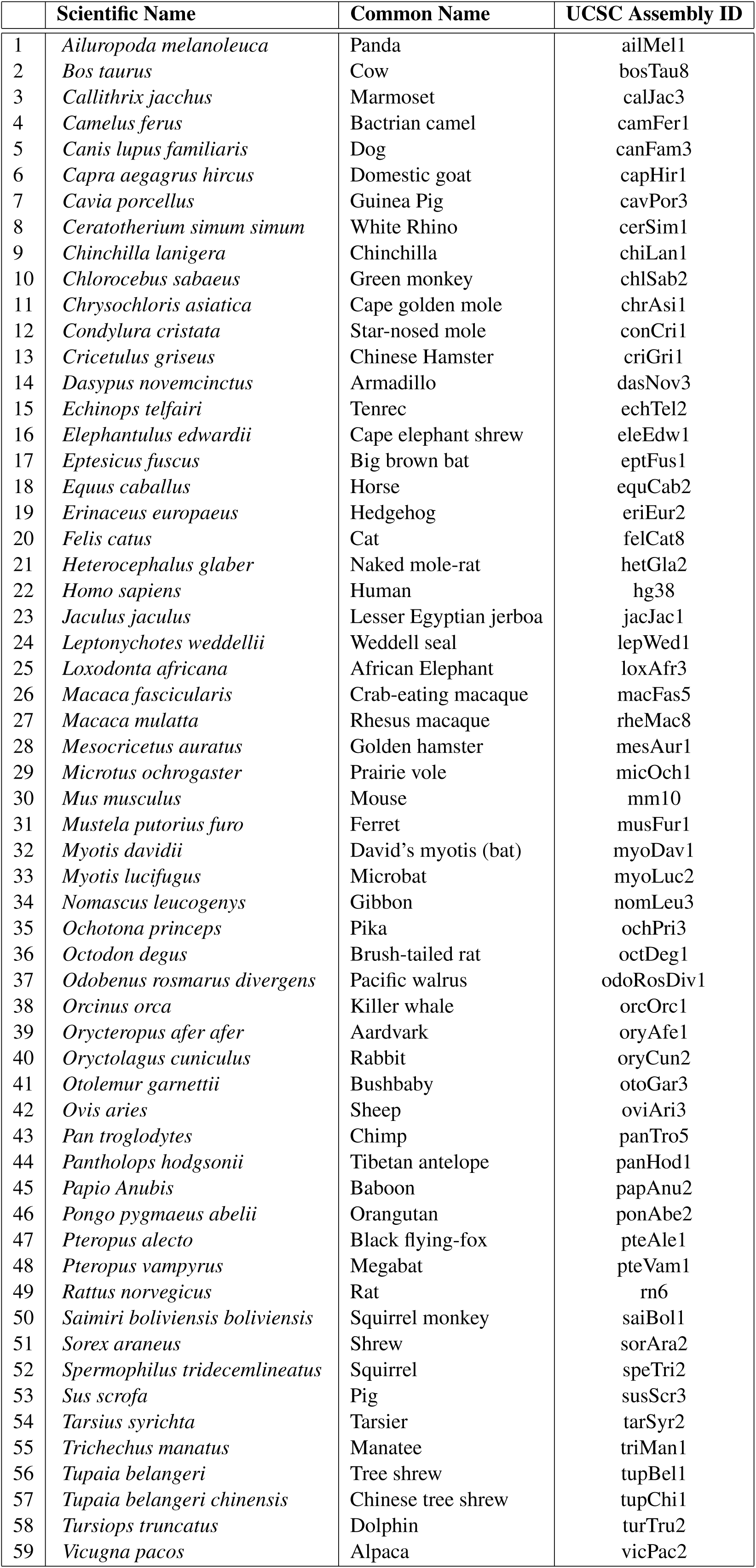
The 59 mammalian genome assemblies used in this study. Scientific name, common name, and the assembly name of the 59 mammalian species (including human) used in this study.

**Supplementary Figure 1.**
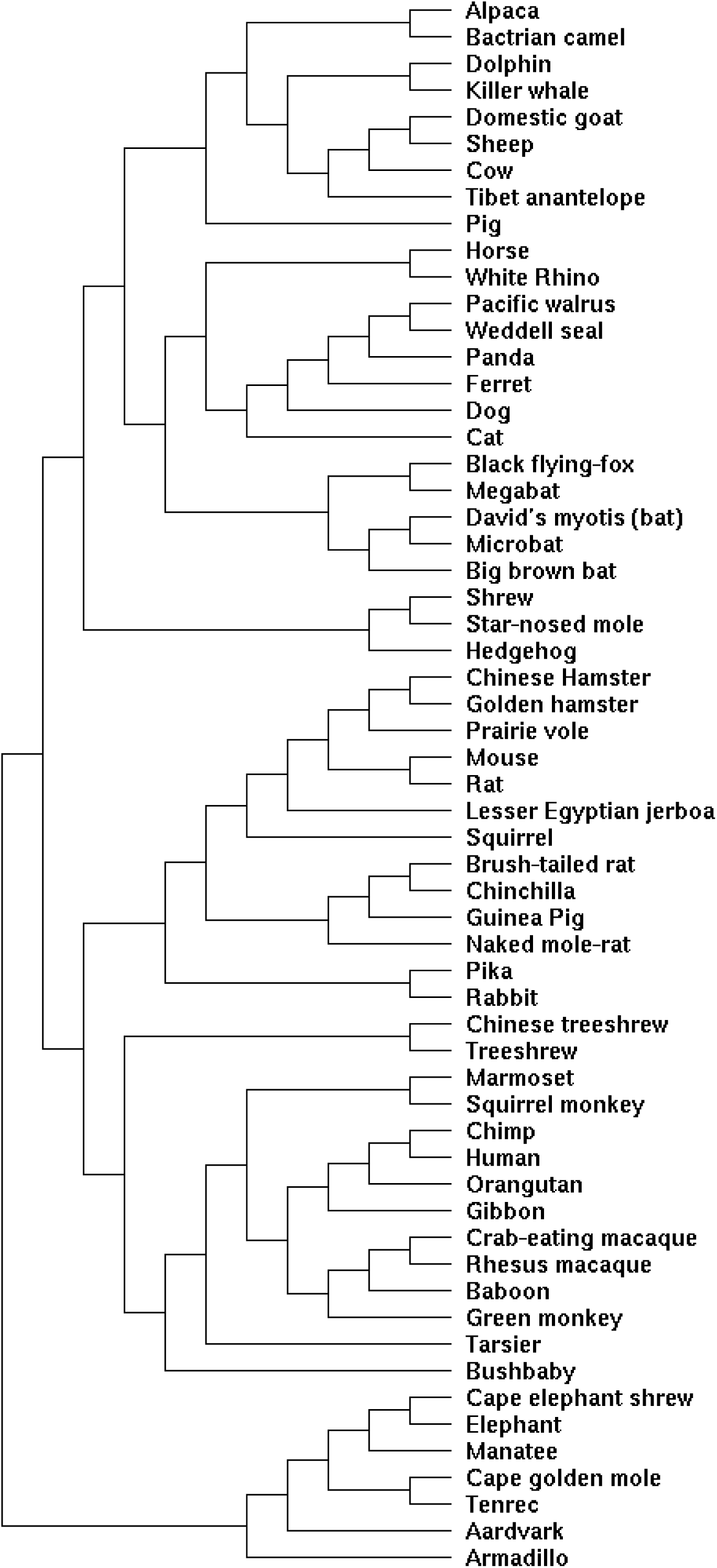
Phylogenetic tree of the 59 mammalian species. The phylogenetic tree of the 59 mammalian species (including human) used in this study. Our pipeline (Figure 3) requires knowledge of only the topology of the tree as shown in this figure and not of the branch lengths.

